# Three-dimensional Cancer Cell Migration Directed by Dual Mechanochemical Guidance

**DOI:** 10.1101/2021.11.11.468299

**Authors:** Pedram Esfahani, Herbert Levine, Mrinmoy Mukherjee, Bo Sun

## Abstract

Directed cell migration guided by external cues plays a central role in many physiological and pathophysiological processes. The microenvironment of cells often simultaneously contains various cues and the motility response of cells to multiplexed guidance is poorly understood. Here we combine experiments and mathematical models to study the three-dimensional migration of breast cancer cells in the presence of both contact guidance and a chemoattractant gradient. We find that the chemotaxis of cells is complicated by the presence of contact guidance as the microstructure of extracellular matrix (ECM) vary spatially. In the presence of dual guidance, the impact of ECM alignment is determined externally by the coherence of ECM fibers, and internally by cell mechanosensing Rho/Rock pathways. When contact guidance is parallel to the chemical gradient, coherent ECM fibers significantly increase the efficiency of chemotaxis. When contact guidance is perpendicular to the chemical gradient, cells exploit the ECM disorder to locate paths for chemotaxis. Our results underscores the importance of fully characterizing the cancer cell microenvironment in order to better understand invasion and metastasis.

## INTRODUCTION

Directed cell migration [1] is of fundamental importance in wound healing [2], immune response [3], and cancer metastasis [4]. In these processes, cells are presented with various types of extracellular cues that bias the direction of their otherwise random motion. Chemotaxis, for instance, is a major type of chemical guidance [5, 6]. By driving a cell to follow the gradient of chemoattractants, chemotaxis provides a unidirectional cue to the cell. Chemotaxis is instrumental in orchestrated functions of multicellular organisms, such as recruiting fibroblasts to acute wound sites [7], and guiding the development of blood vessels [8].

Contact guidance, on the other hand, is a mechanical effect that directs cell morphogenesis and motility based on cues from the topography of 2D substrates [9, 10] or the organization of 3D extracellular matrix [11]. Unlike chemotaxis, contact guidance presents a nematic, rather than a unidirectional, cue to the cells [12]. Migration modulated by contact guidance is observed in T-cells [13], endothelial cells [14], epithelial cells [15], and in the context of cancer [16], where cells directed by contact guidance are also considered to follow a least-resistance path [17, 18].

While the motility of cells under either chemotaxis or contact guidance has been extensively characterized, in realistic physiological conditions chemical cues and mechanical guidance are often present simultaneously. During tumor progression, chemotaxis driven by the gradient of various growth factors and nutrients facilitates the dissemination of cancer cells from primary to metastatic sites [19]. To finish this journey, cancer cells must navigate through the host tissue space filled by fibrous extracellular matrix (ECM) [20]. Here the alignment of ECM fibers is a major source of contact guidance. In fact, it has been shown that the direction of ECM fibers has a significant correlation with tumor prognosis [16, 21].

The migrational response by which a cancer cell integrates multiplexed cues remains poorly understood [22]. The issue is particularly relevant for 3D cell migration, as the disordered microstructure of the ECM has been largely overlooked in the study of cancer cell chemotaxis. In this work, we report a combined experimental and theoretical analysis to elucidate 3D breast cancer cell motility under dual mechanochemical guidance. We show that both ECM microstructure and cell mechanotransduction modulate cancer cell chemotaxis. Despite the complexity of the underlying molecular mechanisms, our results support a simple picture whereby chemotaxis and contact guidance provide additive motility biases to steer 3D cell migration. Taken together, our results underscores the importance of fully characterizing the cancer cell microenvironment in order to better understand invasion and metastasis.

## RESULTS

In order to investigate 3D cancer cell motility in the presence of simultaneous mechanical and chemical guidance, we incorporate flow-induced ECM alignment in 3D chemotaxis assays. In particular, we inject 1 mg/mL neutralized, FITC-labeled type-I collagen solution into the 3D cell migration chamber (Fig. 1A, along the 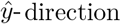). The collagen solution contains low-density RFP-labeled MDA-MB-231 cells, and gradually gelates to form fibrous collagen ECM. See SI for more details.

**FIG. 1.**
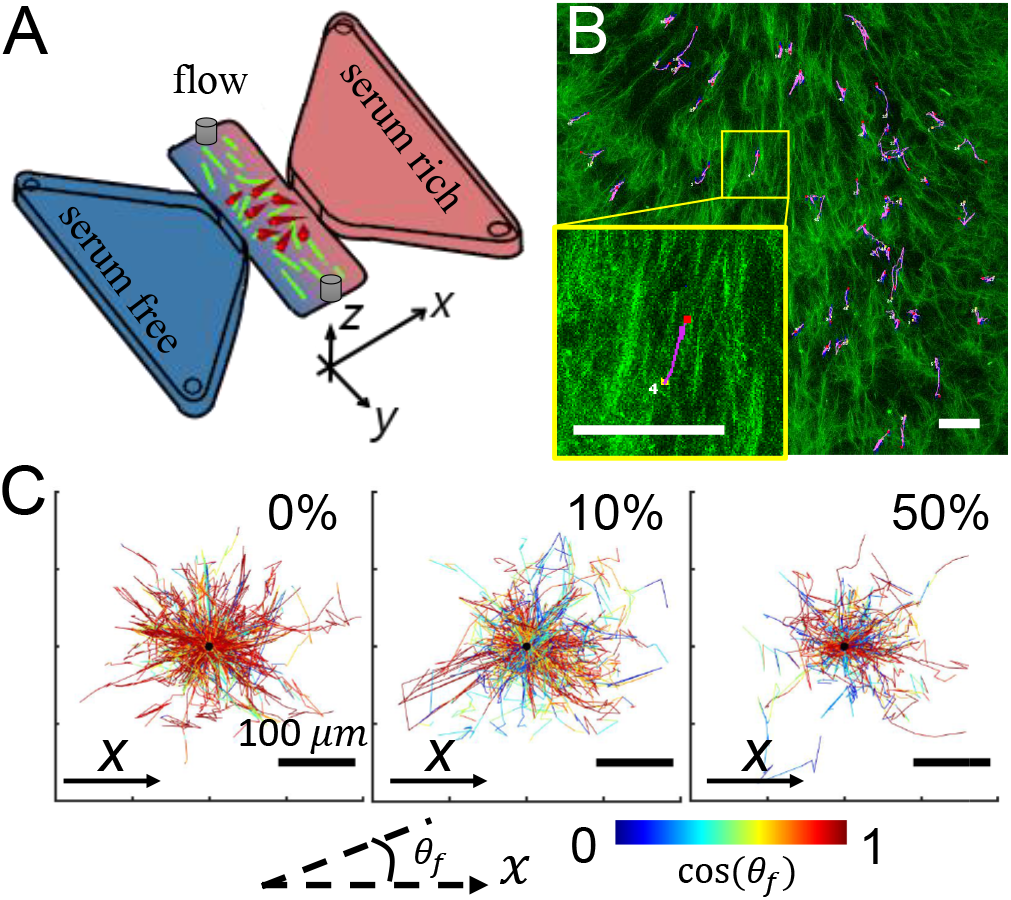
Experimental setup used to investigate 3D cancer cell migration in the presence of dual mechanochemical guidance. (A) Schematics of the dual guidance assay. Chemotaxis driven by a serum gradient is along 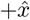 direction. (B) Example cell trajectory (purple) over collagen ECM (green). The yellow box shows a zoomed-in view of a cell following aligned collagen fibers. To compute the direction and coherence of the ECM fiber surrounding a cell, a 150 *μ*m x 150 *μ*m observing window is positioned around the cell of interest. (C) Displacement of cancer cells in the x-y plane under various chemogradient conditions. Here 0%, 10%, and 50% represent the volume concentration of serum in the chemoattractant reservoir, while the other reservoir is filled with serum-free growth medium. The trajectories are imaged every 15 minutes and are colored by the cues agreement cos *θ_f_*. Here *θ_f_* is the acute angle between local ECM direction and 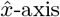. Scale bars: 100 *μ*m.

Using a syringe pump, we program the injection speed and duration, and also alternate the flow direction. The resulting flow field directs the self-assembly of collagen ECM, thereby creating a network of collagen fibers whose geometry vary spatially. Once the ECM is set, we fill the liquid reservoirs with serum-free and serum-rich (10% or 50%volume concentration, 0% as control) growth media. In less than 6 hours, a stable gradient is formed across the migration chamber, which measures 1 mm between the two reservoirs. We conduct live cell confocal imaging of both cells and collagen fibers 6 hours after device setup. Each device is continuously imaged for 12 hours at intervals of 15 minutes, where z-stacks at steps of 10 *μ*m are collected. Because both serum gradient and collagen fiber alignment are primarily in the *x* − *y* plane, we focus on the cell motility in the *x* − *y* projection. Specifically, the chemical gradient drives the cells toward 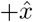 direction, whereas the local mechanical guidance may point along any direction.

The ECM is intrinsically a disordered biopolymer network whose microstructure fluctuates spatially [23]. Therefore instead of bulk characterizations, we must quantify the local contact guidance cues. To this end, we employ image-based methods to measure the principal direction, and coherence (*c*) of collagen fibers in subwindows surrounding individual cells, as reported previously [12] (Fig. 1B). Each subwindow is a cell-centered 150*μ*m×150*μ*m square, cropped from the same z-slice in which the cell is mostly in focus. Coherence *c* varies between 0 and 1, with 0 for completely random aligned fibers and 1 for perfectly parallel fibers. In our experiments, local coherence typically varies between 0.1 to 0.4. The approach of using cell-centered sub-image characterization allows us to compute the instantaneous contact guidance experienced by individual cells.

As a cancer cell navigates the ECM under dual guidance, cos *θ_f_*, where *θ_f_* being the angle between the principal direction of collagen fibers and 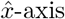, represents the cues agreement. Note that since contact guidance is nematic, we only need to consider the acute angle with 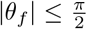. When cos *θ_f_* = 1, contact guidance is parallel with the chemical gradient (ie. along the 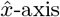). In this case, we expect both cues to work in concert to facilitate chemotactic motion. When cos *θ_f_* = 0, on the other hand, local contact guidance is perpendicular to the chemical gradient. In this case, we expect mechanical and chemical guidance to be in strongest competition for steering the direction of cell migration. In a typical cell trajectory (Fig. 1B) the direction of contact guidance varies along the cell’s migration path. This effect is further demonstrated in Fig. 1C, which shows trajectories (offset by their starting position) of a random subset of cells in different magnitudes of chemical gradient. The trajectories are color-coded by cues agreement cos *θ_f_*. It is evident that while overall a positive chemical gradient biases the cell motility towards serum-rich region, heterogeneous mechanical guidance complicates the task of chemotaxis.

By more closely examining the cell migration in dual guidance, we notice characteristics of the path-finding dynamics. As illustrated in Fig. 2, a cancer cell typically searches for collagen fibers that lie in the direction dictated by chemotaxis. When local coherence is low (Fig. 2A), such fibers are possible to find because collagen fibers are randomly aligned. Conversely, when local fibers primarily align in the direction of chemical gradient (Fig. 2B), cells can readily follow the contact guidance to move towards higher serum concentration. Interestingly, even when collagen fibers are mostly aligned perpendicular to the gradient (Fig. 2C), cancer cells are capable of locating fibers in the direction of chemical guidance. Such fibers are rare in this case, therefore cells spend much time waiting, or moving perpendicular to the chemical cue (i.e. following the mechanical cue) until a path suitable for chemotactic prpgress is discovered.

**FIG. 2.**
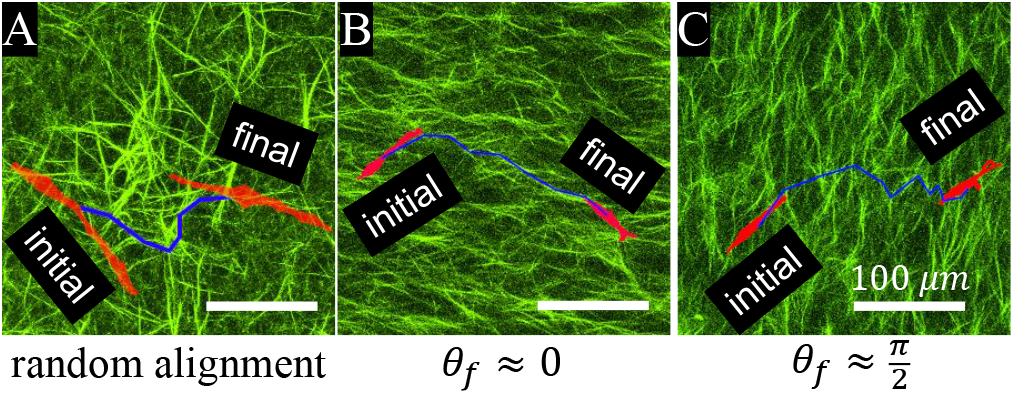
Cancer cell motility integrates dual mechanochemical guidance by exploiting the disordered fibrous structure of the ECM to move up chemogradients. Examples shown here are (A) a cancer cell moves in a region of randomly aligned ECM (low coherence), (B) a cancer cell move along ECM aligned parallel to the chemogradient (*θ_f_* ≈ 0), and (C) a cancer cell takes advantage of ECM disorder to traverse collagen fibers aligned perpendicular to the chemogradient 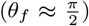. Scale bars: 100 *μ*m. Red: morphology of cells at the initial and final frames. Green: collagen fibers. Blue lines: cell trajectories.

To better quantify these observations, we propose a simple mathematical model to understand how cells integrate simultaneous mechanochemical guidance. In particular, we consider the direction *θ_v_* of cell velocity to follow the Langevin equation, or its equivalent driftdiffusion equation:

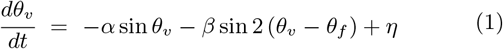

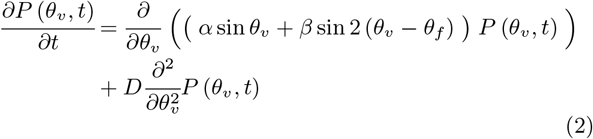

Here *α* > 0 represents the 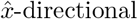 driving by the chemical guidance; *β* > 0 represents the effects of contact guidance; and *η* is a Gaussian random force resulted from both intrinsic and extrinsic noises such that 〈*η*(*t*)*η*(*t′*)〉 = 2*Dδ*(*t* − *t′*). Note that while chemical guidance is unidirectional (invariant under *θ_v_* → *θ_v_* + 2*π*), contact guidance is nematic (invariant under *θ_v_* → *θ_v_* +*π*, or *θ_f_* → *θ_f_* + *π*).

When coherence *c* is low, we expect 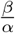 to be small. In this case, cell chemotaxis will navigate randomly aligned fibers regardless of the principal direction of the local ECM. When coherence *c* is higher, the direction of collagen fibers *θ_f_* will have a strong impact of cell motility. Chemotaxis is enhanced when cues agreement is high, as cell path finding becomes easier. Conversely, chemotaxis is curtailed when cues agreement is low, as there are fewer fibers along the chemogradient. Therefore we expect *β* to be positively related with coherence *c*.

We can solve equation (1) to obtain the stationary distribution of *θ_v_*

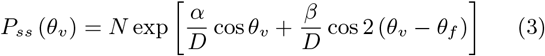

where *N* is a normalization factor. Hence in equation (1), the effects of chemical gradient and contact guidance are represented by two potential energies *V_chem_*(*θ_v_*) = −*α*cos *θ_v_*, and 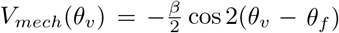 respectively. Therefore our model assumes that cancer cells additively integrate the chemical and mechanical cues, under the impact of intrinsic and extrinsic noises.

To illustrate the model results, we choose generic parameters and consider four distinct scenarios (Fig. 3A). In the absence of a chemogradient 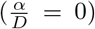, the direction of cell migration is biased solely by the ECM direction. The probability distribution function (PDF hereafter) of *θ_v_* reflects the value of *θ_f_*, and is symmetric with respect to the line *θ_v_* = *θ_f_*. Also in this case *P_ss_*(*θ_v_*) = *P_ss_*(*θ_v_* + *π*), because contact guidance alone is a nematic cue to cell motility.

**FIG. 3.**
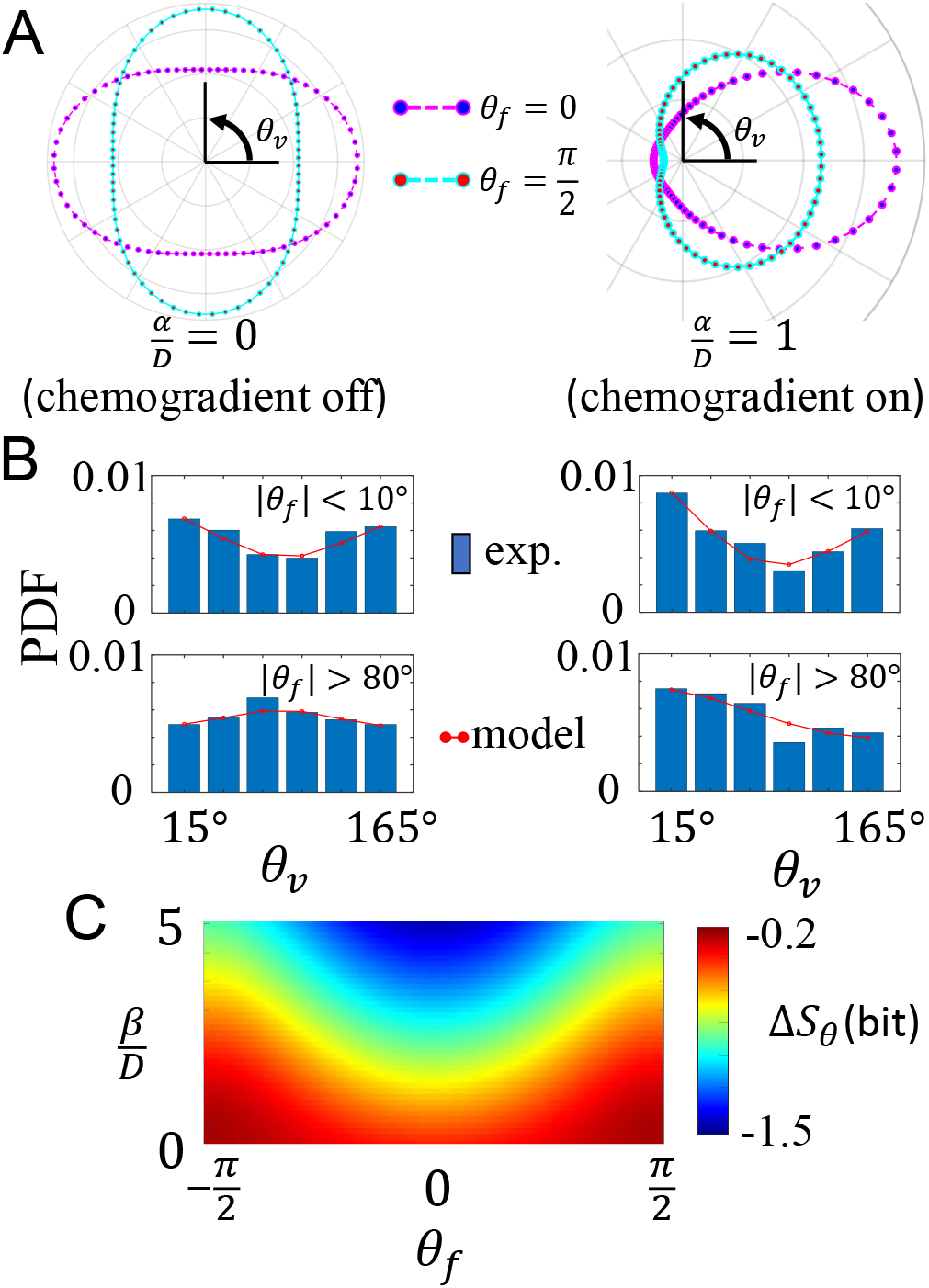
Theoretical model predicts cancer cell motility in response to dual mechanochemical guidance. (A) Probability distribution of cell velocity direction (*θ_v_*) when cues agreement is maximal (*θ_f_* = 0, magenta curve), and when cues agreement is minimal (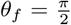, cyan curve). Left: only the mechanical cue is present. Right: both chemical and mechanical cues are present. In these examples we choose 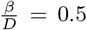 (B) Typical experimental results corresponding to scenarios shown in (A). Abbreviations: PDF: probability distribution function; exp. experiments. For each histogram, *N* >1000 data points are included and the model parameters are obtained using Levenberg-Marquardt algorithm. (C) Uncertainty of cell migration direction quantified by the differential entropy of *θ_v_*. The heat map shows the change of differential entropy Δ*S_θ_* in comparison with the case without external guidance (*α* = *β* = 0). The results in (C) are calculated based on the model by setting 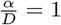.

Once we turn on chemogradient 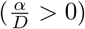, *P_ss_*(*θ_v_*) shifts toward 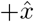 direction. It is informative to consider two special cases with maximal cues agreement *θ_f_* = 0, and minimal cues agreement 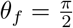. PDF of *θ_v_* is symmetric with respect to *x*-axis, which is again due to the nematic nature of contact guidance. It is evident that chemotaxis index CI, defined by 〈cos *θ_v_*〉, is positive. In general, we note that equation (3) is invariant under reflection symmetry (*θ_f_* → −*θ_f_*, *θ_v_* → −*θ_v_*) and nematic symmetry (*θ_f_* → *θ_f_* +*π*). These symmetries are model-independent.

The four scenarios (with and without chemogradient at maximum or minimum cues agreement), along with the model predicted PDF of cell migration direction, reasonably match with the experimental observation (Fig. 3B). Note that in Fig. 3B we have taken advantage of the symmetry of PDF to map *θ_v_* into the range of 0 to *π*.

The external cues not only determine the average direction to which a cell migrates, but also the uncertainty of its migration direction. To show this, we calculate the differential entropy of *θ_v_* defined as 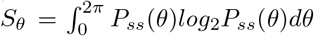. As a reference, the maximum entropy *S_a_* occurs when *θ_v_* distributes isotropically, such that *S_a_* = *log*_2_2*π* in unit of bit.

It is particularly informative to examine the uncertainty introduced by varying mechanoguidance. To this end, we set 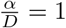, and compute 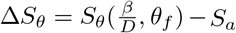 as shown in Fig. 3C. For the range of parameters considered here, external cues reduce the uncertainty of *θ_v_*, such that Δ*S_θ_* < 0. When the strength of mechanochemical cues are fixed (holding 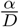 and 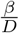 constant), entropy of *S_θ_* is minimized when *θ_f_* = 0, where contact guidance works in concert with chemogradient to reduce the fluctuations of cell migration direction. For fixed chemical guidance, the maximum entropy of *θ_v_* occurs when *α* = *β* and 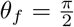. In this case, cells are frustrated by the perpendicular but equally strong cues, thereby maximizing their directional uncertainty.

Our model predicts *β* to be positively correlated with ECM coherence. Additionally, *β* should also increase if cells become more mechanosensitive to contact guidance cues. To test this prediction, we take advantage of the fact that 3D cell contact guidance is primarily regulated by Rho/Rock signaling [11]. In particular, we treat the MDA-MB-231 cells with a well-characterized Rock inhibitor Y27632 and compare the resulting migration directions under 10% serum gradient in collagen matrices. Fig. 4A shows the conditional probability *P*(*θ_v_*|*θ_f_*) for varying angles of the ECM fibers.

**FIG. 4.**
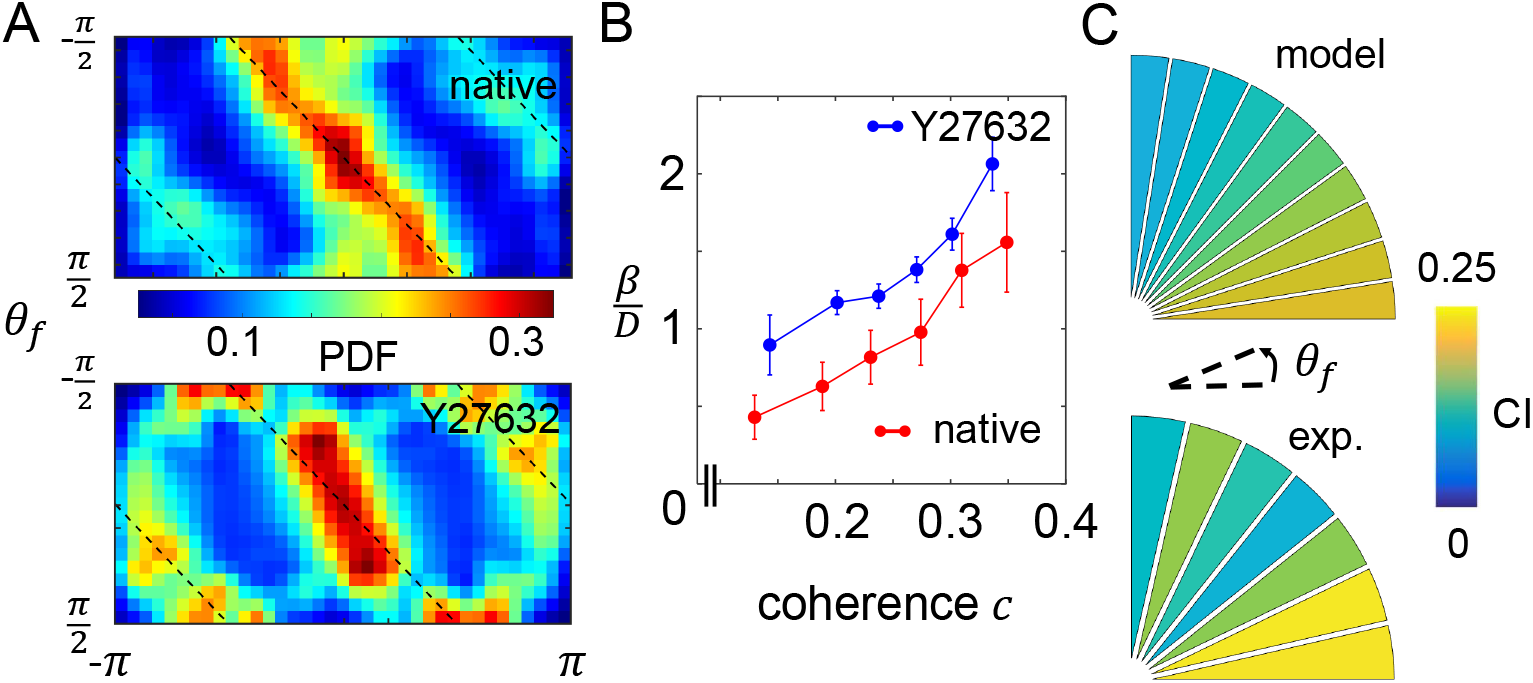
Mechanistic insights into cancer cell motility in response to dual mechanochemical guidance. (A) Experimentally measured conditional probability distribution of *P*(*θ_v_*|*θ_f_*). The data is obtained for cells navigating ECM of relatively high coherence (*c* > 0.2), and with the chemoattractant reservoir filled with 10% serum. A Gaussian kernel has been employed to compute the probability distribution. Data in this figure has also been augmented to satisfy the reflection symmetry: for each data point (*θ_v_*, *θ_f_*), we also include (−*θ_v_*, −*θ_f_*) in the statistics. Top: native MDA-MB-231 cells. Bottom: cells treated with Rock-inhibitor Y27632. (B) At fixed chemogradient (10% across the device), and ECM approximately in parallel to the 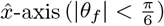, model fitting to experimentally measured probability distribution of *θ_v_* reveals the relation between ECM coherence *c* and model parameter 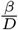. For native MDA-MB-231 cells the best fit yields 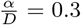 and for Y27632 treated cells, the fitting yields 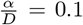. To obtain each fitted data point, a sliding window of *c* with width of 0.1 is used to sample cell velocity at varying ECM coherence. Error bars show the standard deviation of 100 bootstrap results. (C) The chemotactic index 〈cos *θ_v_*〉 for native MDA-MB-231 cells as the direction of mechanical guidance varies. Here we consider the same condition as in (A), where ECM coherence *c* > 0.2 and the chemogradient is 10% across the device. Top: model prediction using fitted parameters as shown in (B). Bottom: experimental measurements. Abbreviations: CI: chemotactic index; exp.: experiment.

For both native and Y27632 cells *P*(*θ_v_*|*θ_f_*) show peaks in the vicinity corresponding to perfect contact guidance (black lines in Fig. 4A). However, the deviations indicate the effect of chemotaxis. In particular, native cells are more responsive to chemical gradient, as the probability near *θ_v_* = 0 is significantly higher than the probability near *θ_v_* = ±*π*. Y27632-treated cells, on the other hand, have more pronounced peaks near *θ_v_* = ±*π*. These observations suggest that Rho-inhibition suppresses cell response to chemical guidance but enhances response to contact guidance. These observations are also consistent with previous reports on the effects of Rho/Rock-inhibition, which promotes the mesenchymal phenotype of cancer cells [24–26], and renders them more sensitive to contact guidance [12].

To further elucidate the effect of Rho/Rock signaling in regulating cell migration under dual guidance, we fit experimentally measured conditional probability *P*(*θ_v_*|*θ_f_* ≈ 0) with equation (3) to obtain quantitative relation between mechanochemical cues and model parameters. The fitting results confirm that while Y27632 treatment reduces cell sensitivity to chemogradient (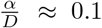 for Y27632 treated cells compared with 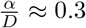 for native cells), Rock-inhibition leads to stronger response to contact guidance (Fig. 4B). These results agree well with our model expectations.

Leveraging the quantitative relation between ECM coherence and model parameter, we then elucidate the impact of ECM principal direction to the efficiency of chemotaxis. To this end, we compute for each fiber direction (within equally spaced binning windows) the mean chemotaxis index *CI* = 〈cos *θ_v_*〉 under 10% serum gradient, and relatively strong coherence (*c* >0.2). Here the average takes into account of experimentally measured distribution of ECM coherence, and therefore reflects the effect of ECM structural disorder. The model predicts, and experiments confirm, that as ECM direction rotates from perpendicular to parallel to the chemogradient, chemotaxis index can increase by as much as two fold.

## DISCUSSION

Tumor metastasis requires cancer cells to navigate through 3D ECM which contains rich mechanochemical cues. In this letter, we investigate the motility of breast cancer cells which simultaneously experience serum gradient and ECM contact guidance, a combination that mimics key aspects of the physiological microenvironment of tumors [27, 28]. We find that over the course of chemotaxis cells experience a contact guidance cue that varies spatially in strength and direction, complicating the task of finding an effective path. The cross-talk of external cues, one that is unidirectional, and one that is nematic, controls the probability distribution of local cell migration direction. As ECM fibers rotate from perpendicular to parallel with chemoattractant gradient, chemotaxis index can increase by more than two fold. We show that in the presence of dual guidance, impact of ECM alignment is determined externally by the coherence of ECM fibers, and internally by cell mechanosensing Rho/Rock pathways. Our results suggests that directed cancer cell migration during metastasis is jointly regulated by biochemical and biophysical cues. Therefore a comprehensive understanding of tumor microenvironment is imperative to predict and control cancer cell invasion.

We note that during the course of metastasis, cancer cell may experience other forms of mechanochemical cues. Many growth factors and cytokines, such as TGF-α and EGF are potent chemoattractants. Other physical aspects of the ECM, such as rigidity and porosity, can also guide cancer cell motility. Interestingly, these mechanochemical cues can be self-generated by cancer cells and their associated stromal cells as they actively remodel the tumor microenvironment [29, 30]. We expect the endogenous cues to coordinate complex collective behaviors, which has just begun to be understood [31].

While our phenomenological model takes into account key aspects that determine the fluctuations of cell migration direction, it can be generalized in several ways. One extension is to include durotaxis effects, as we have explored previously [32]. Dependence on cell morphology [33] could also be included by making the driving coefficients alpha and beta and the diffusion constant dependent on cell aspect ratio. Finally a more mechanistic treatment, such as by explicitly considering the physical interaction of cells with fibrous ECM [34, 35], could directly predict how the effective parameters are determined by both cell and ECM degrees of freedom.

Part of this research was conducted at the Northwest Nanotechnology Infrastructure, a National Nanotechnology Coordinated Infrastructure site at Oregon State University which is supported in part by the National Science Foundation (grant NNCI-1542101) and Oregon State University. P. Esfahani and B. Sun are supported by DOD award W81XWH-20-1-0444 (BC190068). B. Sun is also supported by the National Institute of General Medical Sciences award 1R35GM138179. The work of H. Levine and M. Mukherjee was supported by NSF (PHY-1935762) and NSF (PHY-2019745).

